# Probabilistic Multiple Sequence Alignment using Spatial Transformations

**DOI:** 10.1101/2024.10.12.617969

**Authors:** Sebastián García López, Søren Hauberg, Wouter Boomsma

## Abstract

Multiple Sequence Alignment (MSA) has long been a prominent and critical tool in bioinformatics and computational biology. Its importance lies in its ability to provide valuable insights into the relationships between sequences and the evolutionary pressure leading to amino acid preferences at particular sites in a protein. Despite the recent advances in protein language models, MSAs remain critical in many applications, e.g. for state-of-the-art prediction of 3D structure and protein variant effects. Sequence alignment is typically considered a deterministic preprocessing step, leading to a single static MSA. Especially for low-similarity sequences, parts of an alignment will be subject to substantial uncertainty, which is disregarded when processing a static MSA. Earlier, HMM-based approaches handled this uncertainty by considering the full posterior ensemble over alignments. In this paper, we explore whether a similar approach is feasible within a modern deep learning approach, where we move beyond the Markovian restrictions of earlier models. In particular, we consider whether we can learn the alignment process as distribution over spatial transformations, in combination with a deep latent variable model of protein sequences. A proof-of-concept implementation of this work is available at https://github.com/deltadedirac/Explicit_Disentanglement_Molecules.

## 1 Introduction

For decades multiple sequence alignments (MSAs) have played a central role in the computational modeling of protein sequences and a long list of downstream prediction tasks, ranging from early work on protein secondary structure [1] to the recent breakthroughs in 3D structure prediction [2]. For such downstream applications, the alignment procedure is typically considered a *preprocessing* step, conducted once for a given set of sequences. However, there will generally be errors in the alignments caused by poor sequence coverage or approximations in the alignment algorithms. The effect of such errors on downstream performance is rarely analyzed.

From a probabilistic perspective, it would be desirable to model the *distribution* of possible alignments and “integrate out” the alignment, to take the relevant uncertainty into account. This goal was achieved many years ago in the profileHMM model [3], a hidden Markov model which directly modeled insertions, deletions and substitutions and thus constituted a statistical model over possible alignments, which exactly allowed for averaging over the posterior of alignments [4].

While profileHMMs have been highly impactful, and for many years constituted the de facto standard for describing protein families [5], they are no longer the most effective way to describe the amino acid preferences for individual sites within a family. The Markov-assumption underlying HMMs prevents direct modeling of correlations between sites that are distant in sequence but proximal in 3D space. Models such as Potts models and deep latent variable models (VAEs) have been shown to model such effects more reliably [6, 7]. A disadvantage with these methods is that the alignment process is no longer part of the model, and we lose the ability to probe the sensitivity to uncertainty in the alignments. Therefore, a natural question is whether we can combine the benefits of modern protein family models with a probabilistic description of the alignment process. This is the goal of the current manuscript.

Despite the fact that ProfileHMMs could in principle be trained on raw protein sequences, the corresponding optimization problem during training was known to be difficult, and ProfileHMMs were therefore typically built from pre-aligned sequences in practice [4]. In this manuscript, we will attempt to replicate this approach in the setting of deep latent variable models (VAEs) for protein families. In short, we will provide our models with an initial set of aligned sequences, but attempt to model the alignment process within the model, such that we can align sequences directly to the model, and “integrate out” the alignment by sampling alignments from the posterior.

Our approach is based on the framework of Disentanglement Representation Learning, in which we consider the sequence alignment as an invariant representation that can be disentangled from the raw sequence, and the alignment itself is modeled through a diffeomorphic spatial transformation. In this context, MSA is framed as a parametric spatial transformation problem, where the goal is to infer the optimal transformation by warping the input sequences to achieve the sequence alignment. Since parametric spatial transformations depend on transformation parameters, these can be inferred within a probabilistic graphical model via variational inference.

We summarize our contributions as follows:

- We propose that multiple sequence alignment can be approached as a spatial transformation problem, where the optimal transformation is derived via variational approximations. The concept behind this methodological approach is illustrated in Figure 1.
- We adapt Continuous Piecewise-Affine Based (CPAB) transformations to discrete applications.
- We construct a probabilistic graphical model that allows the generalization of alignments on sequences outside the training set.
- The graphical model enables uncertainty quantification for aligned sequences, which we anticipate can improve performance in downstream tasks.

**Figure 1:**
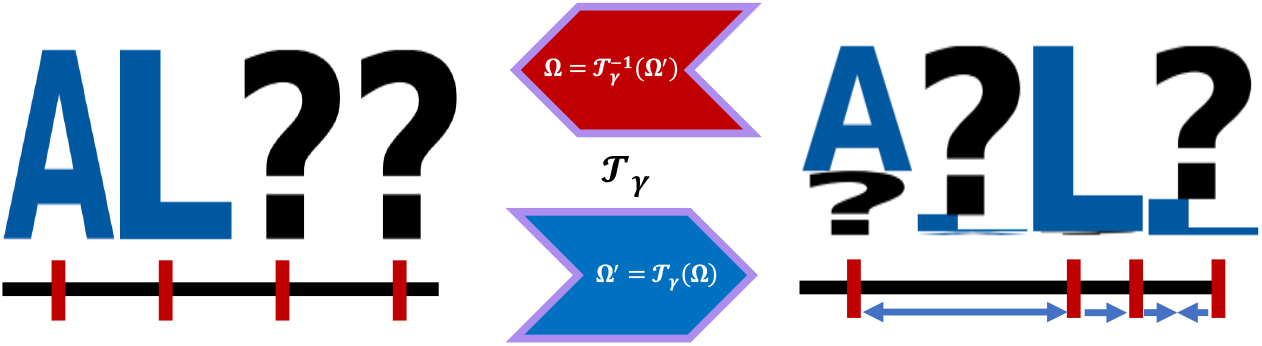
Proposed Methodological Approach: The core idea frames sequence alignment as a spatial transformation problem. Each sequence is represented as an affine grid system, with indices corresponding to individual residues. When passed through the spatial transformer, the system undergoes deformations (indicated by small blue arrows), adjusting the grid and displacing residues to achieve the desired transformation. The optimal estimation of transformation parameters will be addressed using probabilistic modeling, as detailed throughout this work. The transformation, being diffeomorphic, enables the transition between mappings, that is, between the original space of unaligned sequences and their transformed representation of aligned sequences.

## 2 Related work

Algorithms for multiple sequence alignment has been a topic of interest for more than 30 years. Early work established the foundations for efficient calculations of alignments using dynamic programming [8]. Later, hidden Markov models were used to provide a statistical model of protein sequences related by evolution, using discrete latent states to capture insertions, deletions and substitutions [3]. Once trained, these models produce a multiple sequence alignment of a set of sequences using a dynamic programming algorithm. These so-called Profile HMMs became a standard technique for describing protein families [5], and for homology search [4].

While Profile HMMs accurately capture the amino acid propensities at each site of a protein, their sequential nature makes them ill-suited for describing non-local correlations in a protein sequence. In 2011, several works demonstrated that such pairwise effects could be efficiently modeled using Potts models [6, 9], and that these correlations provided important signals for 3D structure prediction [9]. Higher-order effects were too numerous to model efficiently with the same technique, but later work showed that such correlations could be captured through the continuous latent variable of a variational autoencoder [7]. The likelihood of such models was later shown to correlate well with the pathology of clinical variants [10], and other variant effects [11].

In the last few years, protein language models have emerged as powerful tools for protein analysis. While such models are potential alternatives to multiple sequence alignments, there are still many cases where they are outperformed by alignment-based methods, especially when many homologous sequences are available. As a consequence, hybrids of the two modeling approaches have been proposed to obtain more robust performance [12]. Other approaches have modelled the alignments themselves using language-model like approaches [13]. Finally, recent work has provided differentiable implementations of the alignment procedure, making it possible to differentiate through the alignment for downstream predictions [14].

Closest to our work is that of Weinstein and Marks [15], who propose a structured observation model that avoids the need for preprocessing sequences into an alignment, and was shown to formally generalize earlier work such as the profile HMM. Our work aims to provide a more efficient alternative by directly predicting the transformations rather than inferring them through a dynamic programming approach.

## 3 Methods

Our basic approach is illustrated in Figure 2. The purple regions represent the random variables, including latent variables, while *x* and 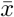 denote the input and output amino acid sequences, respectively. In Figure 2.C,D, the right branch of the models is a variational autoencoder (VAE) of aligned protein sequences, similar to the DeepSequence model [7], while the left branch is a variational autoencoder that outputs parameters for a spatial transformation. The goal is to infer the optimal transformation 𝒯_*γ*_ that allows enough deformation to spatially shift the residues to the proper alignment. Spatial transformations are parametric because they rely on transformation parameters (Section, and these parameters must be inferred (variationally) by inserting them into a graphical model as latent variables. Previous work has successfully inferred the parameters of diffeomorphic spatial transformations to capture invariant representation using graphical models. The model from Detlefsen et al. [16] that inspired our proposal, shown in Figure 2.B, is known as the Conditional Variationally Inferred Transformational Autoencoder (C-VITAE).

**Figure 2:**
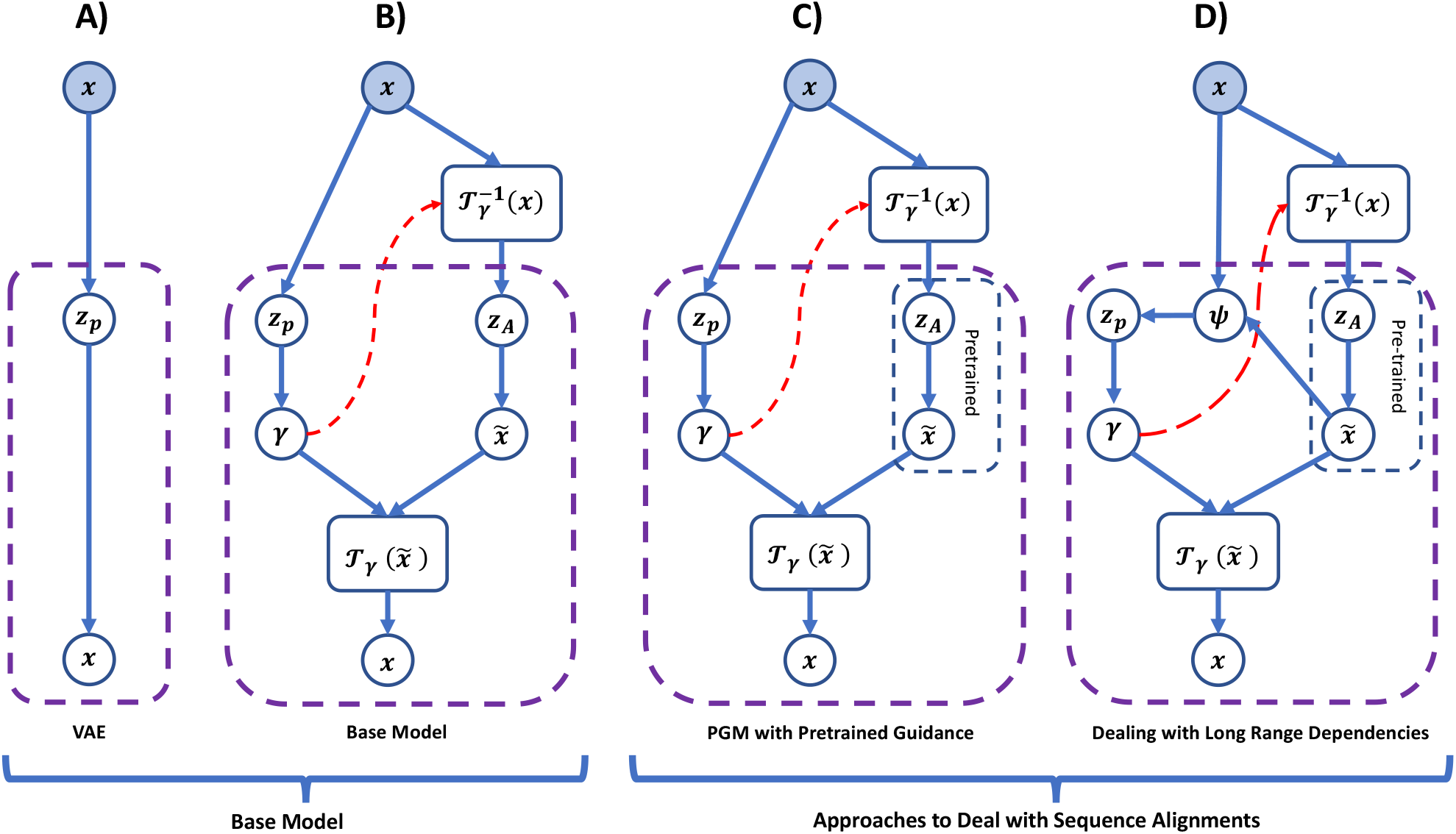
Proposed Framework: **A)** shows the graphical representation of a Variational Autoencoder (VAE) model, while **B)** shows the graphical representation of the Conditional Variationally Inferred Transformational Autoencoder (C-VITAE) [16]. These models have served as reference frameworks, inspiring the development of this work. On the right, **C)** illustrates the adaptation of the method from [16] for MSA and **D)** shows the model adaptation to deal with long-range dependencies. The proposed strategy partially adapts the methodology from [16] to demonstrate that MSA can be approached as a spatial transformation problem. Key distinctions from the original work in Part B include: 1) adaptation of the CPAB transformation, denoted as *T*_*γ*_, to handle categorical distributions (see Section [17]); 2) incorporation of a pretrained block, indicated by blue dashed lines, to guide the alignment process as an informative prior; and 3) integration of a graphical model that introduces a new latent variable (Ψ) aimed at capturing more accurate features, thereby mitigating flat optimization landscapes.

As the name suggests, C-VITAE is a generative model structured around a variational autoencoder framework (highlighted in purple in Figure 2.B), involving two latent variables, *z*_*P*_ and *z*_*A*_. One of these variables, *z*_*P*_, parameterizes the transformation parameters, *γ*, which induces the transformation 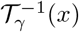. This transformation captures the invariant representation in *z*_*A*_, facilitating the disentanglement of unique sample attributes encoded in *z*_*P*_, as well as the invariant features represented in *z*_*A*_. The full model can be viewed as a large variational autoencoder, where the reconstruction of the original samples *x* is achieved through 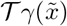, using the same *γ* parameters computed for both 𝒯*γ*^−1^ and 𝒯_*γ*_. In practice, the graphical model is composed of 2 VAEs, i.e. the first one going from *z*_*P*_ to *γ* and the second one going from *z*_*A*_ to 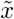, with two layers of spatial transformations (𝒯 *γ*^−1^ and 𝒯_*γ*_ respectively) sharing the same transformation parameter *γ*, which is inferred during training. This is one of the reasons why a fully diffeomorphic spatial transformation is required for modelling purposes, as it allows a return to the observation space for the reconstruction measurements necessary for the computation of the variational approximation. We use the Continuous Piecewise Affine-Based Transformations (CPAB) as a parametric spatial transformation (see [18] and Sections 3.1).

In the original work (see [16] and Figure 2.B), the graphical model evaluated its generative and representational capabilities on image datasets such as MNIST, SMPL, and CelebA, which were modeled as continuous variables. Since protein sequences are inherently categorical and spaced in discrete intervals, we cannot apply the original graphical model directly. Furthermore, we found that the original C-VITAE model struggled with longer-range dependencies. To address these limitations, we propose the schemes shown in Figure 2.C and Figure 2.D. The models use a pre-trained variational autoencoder (VAE) on an initial set of aligned sequences as a prior to guide the alignment process, and introduces a Gaussian process based smoothing approach to model the discrete signal (see section 3). In addition, we propose another graphical model (right side of figure 2.D) that incorporates an additional latent variable to get a richer featurization to achieve more expressive transformations, thus mitigating the problem of long-range dependencies. The corresponding mathematical derivations for its evidence lower bound (ELBO) appear in sections 3.1.3 and 3.1.4.

### 3.1 Spatial Transformation Layers

As our goal is to define multiple sequence alignment as a spatial transformation problem, it is crucial to first introduce a notion of what a spatial transformation is. Spatial transformation techniques have long been fundamental to image processing and computer vision, allowing modifications on the input space, i.e. rotation, scaling and translation with the purpose of achieving invariance in the representation space. This ensures that the representations remain stable under various transformations or changes in the input, thereby enhancing robustness and generalisation [19], [17].

Spatial Transformer Networks (STN) [17] parameterize a neural network as a localisation network that processes the raw input to extract feature maps that will work as transformation parameters. These transformation parameters are fed into a grid generator which performs an affine transformation to warp a regular uniform grid given the transformation parameters estimated by the localization net. The output from the grid generator, along with the original input, is used by a sampler to perform interpolation to produce the final transformed output. The attractiveness of STN lies in their ability to apply flexible, non-rigid deformations to signals, thereby enabling the extraction of invariant features for representation. This makes them useful in tasks such as image registration, among other applications [17].

For our purposes, we require invertible spatial transformations. We therefore adopt on a special type of spatial transformation method called Continuous PiecewiseAffine Based Transformations (CPAB) [18]. CPAB is a transformation that allows the parameterisation of non-rigid, smooth and differentiable deformations in the input space, while retaining the property of being fully diffeomorphic. The attractiveness of CPAB lies in its ability to provide highly expressive transformations at low computational cost, making it well suited to probabilistic modelling techniques such as variational inference, Markov Chain Monte Carlo, among others [16], [18]. CPAB has also been successfully incorporated as a structural element to enhance the expressiveness of existing STNs. For instance, Diffeomorphic Transformer Networks [20] replace traditional affine transformations with CPAB, improving expressiveness and performance in classification and regression tasks and the Probabilistic Spatial Transformer Networks [21] which provide a stochastic extension of conventional STNs. Since CPAB is a parametric transformation, the optimal transformation parameters can be estimated as a natural part of the parameters in the variational autoencoder.

#### 3.1.1 CPAB Transformation — Fundamentals

Continuous Piecewise-Affine Based Transformations (CPAB) [18] are inspired by conventional affine transformations. Affine transformations are mathematical functions that allow mapping in a way that preserves the spatial representation under geometric changes such as scaling, rotation and translation. Despite their widespread use in image processing, affine transformations are limited by their linear nature, making them unsuitable for modelling non-linear deformations due to their inability to capture complex spatial relationships [22], [17]. Beneficially, affine transformations are, however, diffeomorphic and differentiable over their entire domain.

To overcome the rigidity of conventional affine transformations, the input space is partitioned into multiple regions and different affine transformations are applied to each region. This approach is also known as Continuous Piecewise Affine Transformations (CPA). While CPA increases flexibility, it introduces non-differentiability at the region boundaries, resulting in smooth transitions within regions but not across boundaries [18]. CPAB introduces two changes compared to the CPA approach. First, linear constraints are applied to ensure both continuity and differentiability across the entire transformation space, effectively linking each partitioned region to maintain global continuity. Second, the deformations are produced through the integration of CPA vector fields [18].

The CPAB transformation process starts by partitioning space into small regions using tessellation cells. This creates a grid system that is used in the deformation process. Then, affine functions are applied to construct CPA vector fields for each region, and linear constraints are imposed to ensure continuity across the regions. Each associated affine matrix generates a vector field that facilitates mapping of the initial point to its new position using the transformation parameters *𝓋*^*ζ*^. Finally, the integration of trajectories generated by the vector fields, followed by interpolation, yields the final transformed output produced by CPAB [18, 21]. The transformation is defined by the trajectory of the vector fields, which is determined by solving the following differential equation:

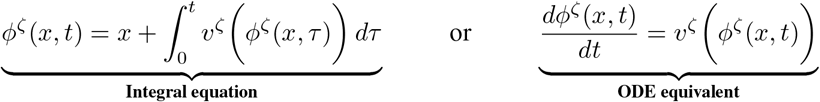

where *v*^*ζ*^ represents the transformed grid point within the set of affine matrices 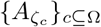 and Ω is the transformation domain [18]. These transformations define the diffeomorphic map *T*^*ζ*^ : *x* ↦ *ϕ*^*ζ*^(*x*, 1).

#### 3.1.2 CPAB Transformation — Adaptation to Discrete Domain

The standard implementation of CPAB transformations relies on linear interpolation between the deformed affine grid, adjusted by the transformation parameters, and the initial input space to produce the final transformed output. While this approach efficiently transforms continuous signals, it faces challenges when quantifying uncertainty within discrete states. Fortunately, the interpolation in CPAB is decoupled from the rest of the components that enable the affine grid deformations. Since the deformations on the affine grids take place first (the transformation component), to then translate these changes into the output space by mapping, we can think of making these transitions by probabilistic means.

In the context of proteins, each sequence can be viewed as a set of categorical distributions, where the highest probability is assigned to a given residue occurring. Thus, each protein can be represented as a sequence of one-hot encoding vectors. We convert this into a continuous representation by placing the residue observations to occur at unit intervals on the real line, and applying a smoothing operation. For our purpose, we require a smoothing operation, that distributes probability mass locally around the discrete observations, gradually dropping off to a prior probability corresponding to a uniform distribution of the 20 amino acids. We employ Gaussian Processes (GP) for this purpose. Formally, this would require a multi-output GP (MO-GP), that takes into account that the probabilities for the twenty amino acids are coupled by a requirement to sum to one. However, for simplicity, we model the development for each amino acid propensity independently, and renormalize post-hoc. The strategy is visualized in Figure 3.

**Figure 3:**
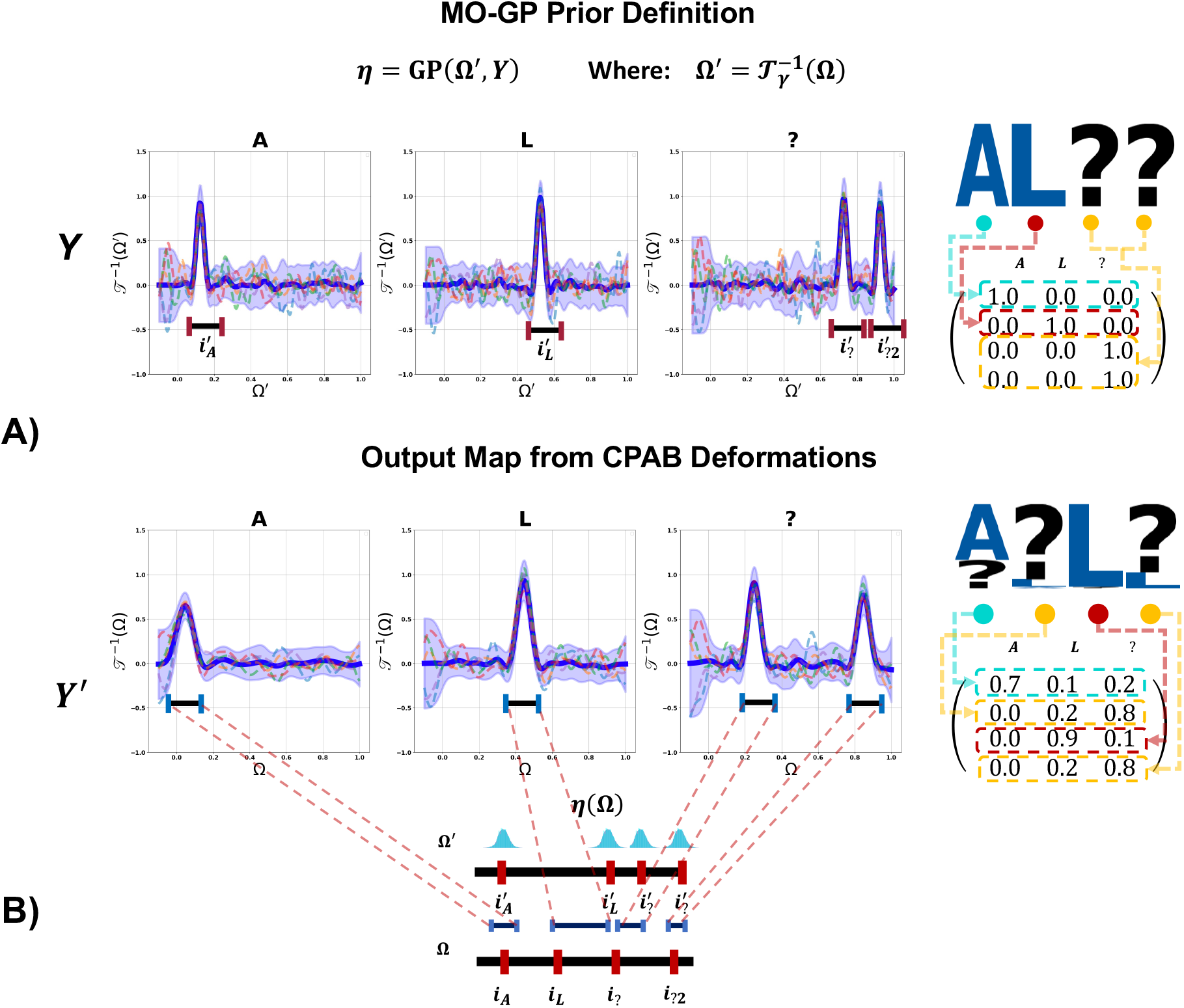
Side A illustrates the preparation of the MO-GP prior, where the deformed vertices of the affine grid Ω′, denoted by **i**, are generated by the CPAB transformation. These serve as input values, while the output values correspond to the one-hot representation *Y* of the original input sequence. Side B shows how the output mapping would be performed. Both the deformed affine grid with CPAB and the uniform affine grid are required to determine the displacement relative to a reference. This allows interpolation via kriging, resulting in the deformed output space *Y*′.

The interpolation method with MO-GP is defined as follows. The index space is composed of two different grid types: one modified by CPAB deformations and the other uniform. An index is assigned to each vertex that defines a part of the tessellation cell. Each index is associated with a residue and its position in the protein, ensuring that the order of the amino acids in the sequence is preserved. Likewise, each index is assigned a one-hot encoding for the corresponding residue, based on a predefined alphabet. The MO-GP prior is defined using the index space Ω′ corresponding to the grids deformed by CPAB and linked to their respective one-hot encoded values *Y*, as shown in Figure 3 (side A). After defining the MO-GP, an affine uniform lattice is used to map the transformation to the output space. The shift relative to the information present in the MO-GP prior is determined by kriging (Figure 3 - side B). It is important to emphasize that MO-GP operates strictly as an interpolator, with no training, optimization or parameter inclusion in further training schemes. However, the choice of an appropriate length scale is crucial as it governs the influence of nearby data points. Specifically, the length scale defines the distance within the index space at which function values are no longer correlated, thus determining the strength of correlation between data points as a function of their distance.

#### 3.1.3 Derivation of ELBO of Basic Framework from Base Model

The initial inference of optimal CPAB transformation parameters for alignment induction and density estimation is based on the Conditional Variationally Inferred Transformational Autoencoders (C-VITAE) [16] (see Figure 2.B). We derive the evidence lower bound by considering the likelihood

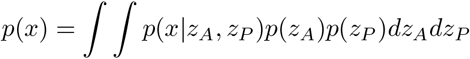

and defining approximate posteriors for the two latent variables

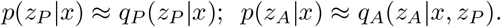

We can derive the marginal likelihood:

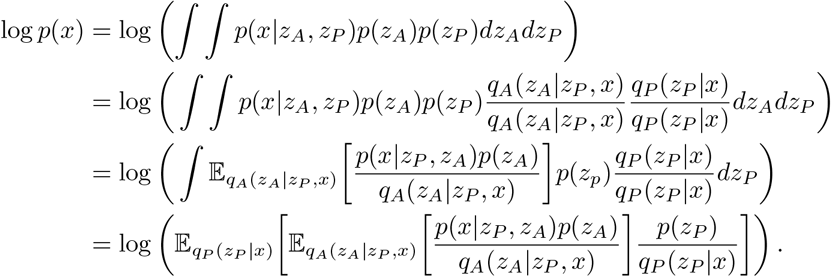

Using Jensen’s inequality to change the order of the external expectation with the logarithm gives

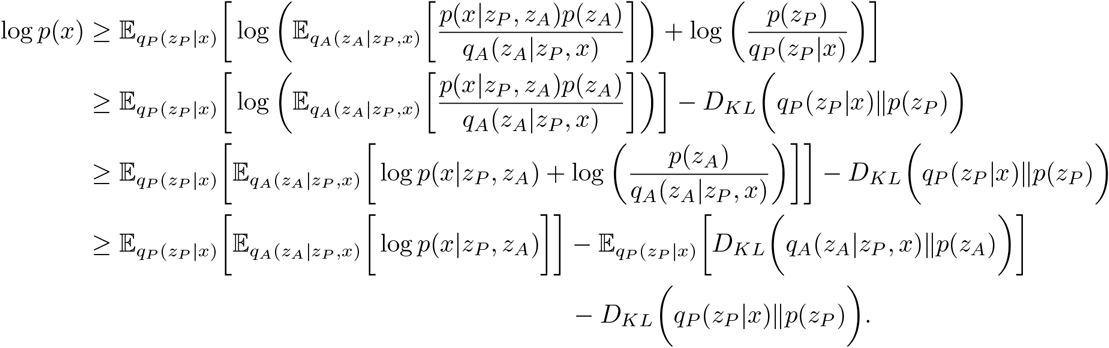

The Evidence Lower Bound (ELBO) can now be written as

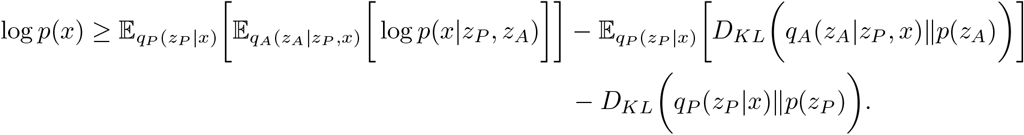

This expression represents the ELBO introduced in [16], which is used as a loss function to optimise the density and obtain the optimal values for the adapted CPAB transformation. This ELBO will also serve as a reference for the derivation of an ELBO specifically designed to address long-range dependencies (see Figure 2.D and Section 3.1.4).

#### 3.1.4 ELBO Derivation from the Model to Deal with Long Range Dependencies with Prior

The primary limitation of the base model (Figure 2.B) is its susceptibility to converge in local minima, particularly when dealing with long-range dependencies in large sequences. This likely arises from an insufficient capacity to capture node-specific information in the graphical model, leading to flat optimization landscapes. To mitigate this, we introduce an additional latent variable as a feature extractor, increasing the expressiveness of *z*_*P*_ and refining the CPAB transformation parameters. The updated model (Figure 2.D) includes a new latent variable, Ψ, which uses samples from *z*_*A*_ to inform the distribution over the transformation latent space *z*_*p*_.

Ψ depends on *z*_*A*_ through 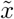, and we can thus write its conditional probability as

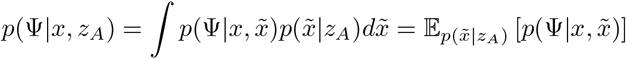

Although this expectation will generally be expensive to evaluate, we only require a rough estimate, and approximate the expectation with a single Monte Carlo sample. In the following, we choose 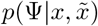 to be a Gaussian parameterized by a light attention block.

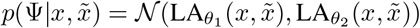

In practice, we found it to work well to set the variance of this distribution to zero, thus effectively using a delta function.

Since *Z*_*A*_ belongs to the prior distribution and remains independent, that is, it is not affected by other latent variables in the graphical model, the model parameters associated with this prior (blue dashed region of Figure 2.D) are fixed. The graphical model thus gives rise to the following factorization

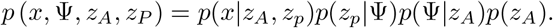

Likewise, the evidence can be defined as

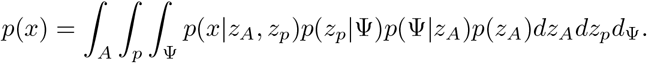

To derive log *p*(x), we get the following

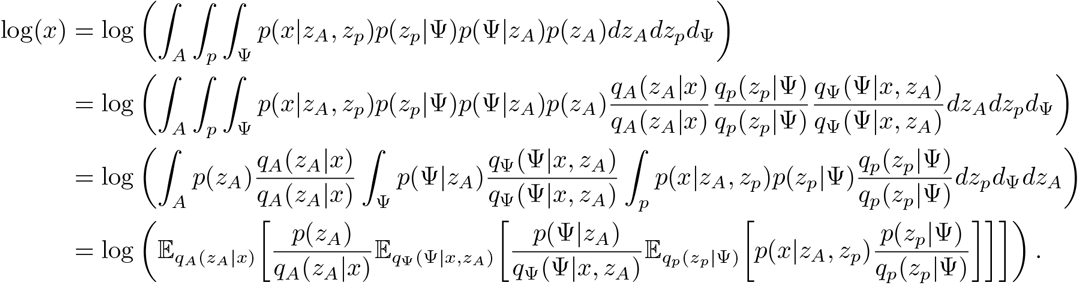

By using Jensen inequality, we get

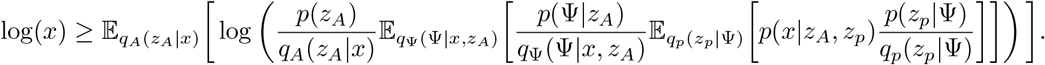

At this point, a small substitution can be introduced to facilitate the algebraic manipulation required to derive the final expression, as follows

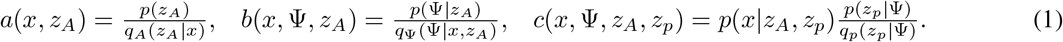

Then, we get

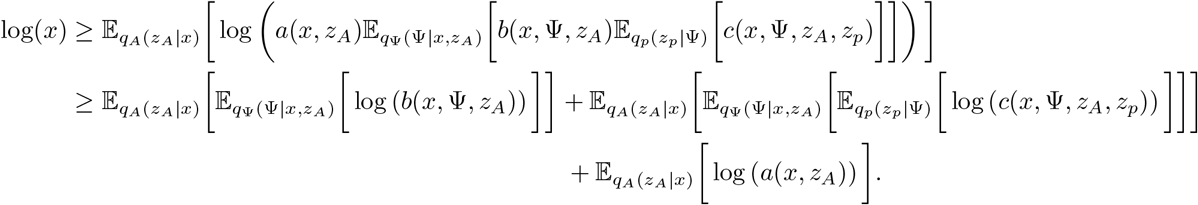

where

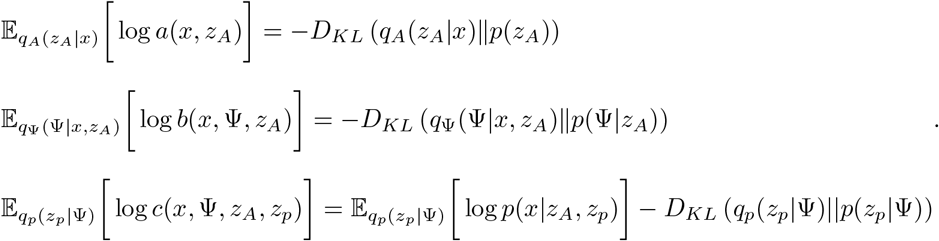

Expanding *a* (·), *b*(·), *c*(·), we obtain our final expression:

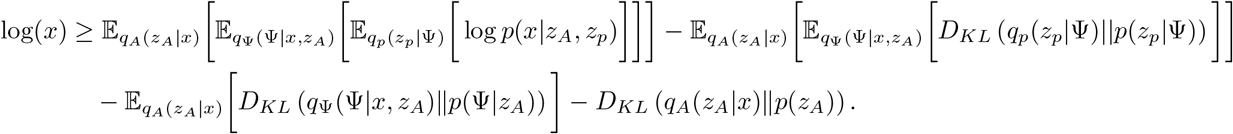

This last expression represents the ELBO that will be used to make the density estimation in the presence of long-range dependencies on Figure 2.D

## 4 Experimental Setup

### 4.1 Experimental Design

To evaluate the effectiveness of our proof-of-concept framework for sequence alignment, two types of experiments were conducted.

#### Experiment 1 - MSA Feasibility on Synthetic Data

The primary objective of this experiment is to assess whether spatial transformations, using the modified CPAB transformations for discrete domains, are expressive and robust enough to produce consistent sequence alignments. To this end, a pre-trained model is built using a Variational Autoencoder with four synthetic sequences, serving as an informative prior to guide the sequence alignment process. The same sequences used to pre-train the prior are also used to train the graphical model, to infer the optimal transformation parameters to achieve the best alignment. The goal is to determine whether, by (over)fitting the density on these sequences, the model can converge to an appropriate alignment and demonstrate robustness as a method for aligning real protein sequences.

#### Experiment 2 - Capacity of Transformations to Generalize Alignments to New Sequences

The second experiment aims to demonstrate that training our model using a pre-trained prior with a very small amount of proteins, can effectively infer the alignment of both sequences employed for density estimation and new sequences to be aligned. It is important to emphasize that the proteins used to train the prior, those for density estimation, and the test set are entirely different, with no overlap among these three groups. Another notable distinction in this second experiment is the use of a limited amount of data to induce the pre-trained prior as well as for density estimation (see Sec. 4.2). The intention behind using relatively small training sets for both the pre-trained and graphical models is to empirically demonstrate the effectiveness of this approach in performing alignments and its ability to generalize such alignments to new sequences beyond the base knowledge

### 4.2 Datasets

For the first experiment, a small synthetic dataset was created, consisting of four synthetic proteins based on a five-character alphabet, including three amino acids and two gap symbols. The pre-trained model used the gap symbol −, while the symbol ? was used for padding in the density estimation model. The rationale behind this is to prevent flat landscapes during the evaluation of the loss function. The same gap scheme has been applied for both experiments 1 and 2. For the second experiment, we used real protein data, specifically those associated with the WW domain (RSP5 domain) [23]. Protein sequences were obtained from InterPro, with manually curated seed alignments. From this dataset, 224 proteins were selected as a pre-trained prior, while 60 proteins were used to train the graphical model. An additional 15 proteins formed the test set. The 224 proteins were previously aligned using Clustal Omega [24], and the pre-trained prior was trained on aligned sequences using variational autoencoders according to the DeepSequence methodology [7].

### 4.3 Details about architectures and parameterization of the model

The construction of the graphical models is based on a composition of interconnected VAEs designed for density estimation [25]. The base model, following [16], consists of a pre-trained VAE model from *Z*_*A*_ to 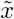, and a block from *Z*_*p*_ to *γ*, which estimates transformation parameters for the spatial transform layers (Figure 2.C). The VAE configuration consists of 3 layers for both the encoder and decoder. The output generated by *γ* in the graphical model varies depending on the tessellation cells selected by the spatial transformation layer, with an 8-cell tessellation used for the first experiment and 1750 cells for the second. A higher number of partitions improves the integration region in the CPAB vector field. Additionally, a length scale of 0.5 was applied to the MO-GP to capture covariance between grid components more effectively, enabling more flexible data transformations. For the light attention layer represented by Ψ (Figure 2.D) in the second experiment, the parameterization follows [26], except for modifications in the input structure, as described in Section A.2. Furthermore, the number of channels in the light attention layer was set to match the alphabet used to represent sequences, i.e., 22 channels.

### 4.4 Optimization Setup

AdamW was used as the main optimiser. The learning rate (lr) was adjusted according to the type of experiment: In the first experiment, the pre-trained model used 1000 epochs with a lr = 1 × 10^−5^, while the density was trained with a lr = 1 × 10^−4^ for 600 epochs. In the second experiment, the pre-trained model was trained for 2000 epochs with the same lr = 1 × 10^−5^, and the density was again trained with a lr = 1 × 10^−4^ for 600 epochs.

## 5 Results and Discussion

### 5.1 MSA Feasibility on Synthetic Data

Figure 4 illustrates the results of the first experiment. A closer examination of the sequence alignment generated (Figure 4.B) reveals that the synthetic amino acids were correctly aligned across columns, indicating that the transformation inferred by the graphical model is robust enough to estimate the transformation parameters accurately and induce proper alignments. These results suggest several findings regarding the use of the adapted transformation 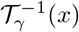 in discrete domains, among them: we demonstrate that the approach based on spatial transformations can indeed successfully align sets of biological sequences in batches. This approach differs from traditional methods based on Markov assumptions, such as HMMs, in that it does not assume alignment based on previous states of the sequences to align residues, but instead uses vertex shifts between residue positions in aligned columns. In addition, it provides the ability to quantify uncertainty in alignments using MO-GP mappings built into the CPAB modification.

**Figure 4:**
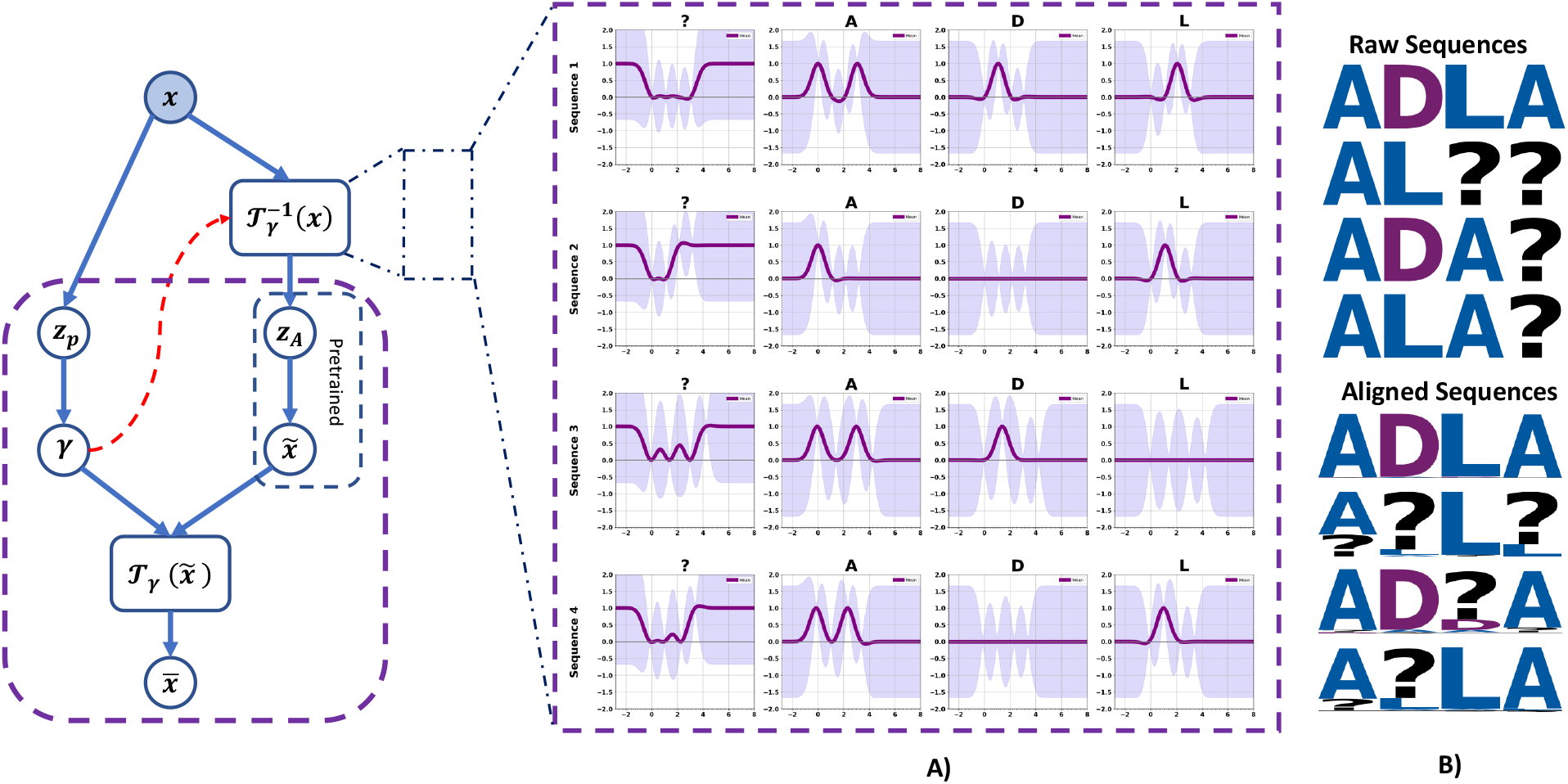
Results of alignments on synthetic data: Side **A)** describes four sequences, each accompanied by their designated names, arranged in rows, and four channels identified by alphabetical symbols. In the GP interpolation plots for each channel, the x-axis represents the index space containing the continuous values of the affine grid and their offset, while the y-axis indicates the probability value of the given state or channel component. In particular, the Multi-Output Gaussian Process (MO-GP) includes four channels or states, represented by the symbols ?, A, D and L, which form the alphabet used for interpolation between discrete states. In addition, the MO-GP prediction used to fit the CPAB transform to the modified CPAB sampler is the mean of the posterior distribution over the channel, with the uncertainty of the prediction represented by a fluctuation region (shown as a purple-shaded area). Side **B)** illustrates the final transformation inferred by the graphical model, where the upper part shows the initial input or its original appearance, while the lower part shows the final result of the sequence alignment as inferred by the spatial transformation layer 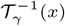.

On the other hand, Figure 4.A shows the interpolations by the MO-GPs within the CPAB modification to estimate the likelihood of each residue given its position in the deformed affine grids. An important observation in the deformed affine grids is that the peaks align with the axes corresponding to the vertices. This alignment is significant because it allows for a geometric interpretation: within each sequence per channel, there should be a positional point of convergence. For instance, if a given column contains A or D, these elements should converge spatially to a similar point along the transformed grids for each sequence. In general, the Spatial Transformer is expressive enough to perform probabilistic alignments consistently. However, it must be acknowledged that this evaluation is based on an overfitting of the graphical model using identical sequences for both input and training of the pre-trained model (aligned sequences for the prior and unaligned for global model). It remains to be determined whether this observed performance would be maintained if the sequences used for pre-training the model were completely different from those used for training the graphical model, as well as for the new sequences to be evaluated given the trained density. In addition, the ability of the model to generalise the alignment to novel sequences is an important consideration. These issues are explored in the next section.

### 5.2 Capacity of MSA generalization to new sequences

The second experiment use real-world data from the WW domain (Figure 5) to assess the ability of the graphical model to generalise sequence alignments for novel sequences and to deal with long-range dependencies in large sequences (see Figure 2.D). In particular, none of the protein sets used for pretraining, density estimation, or testing had any common protein, ensuring robust alignment for a completely novel sequence.

**Figure 5:**
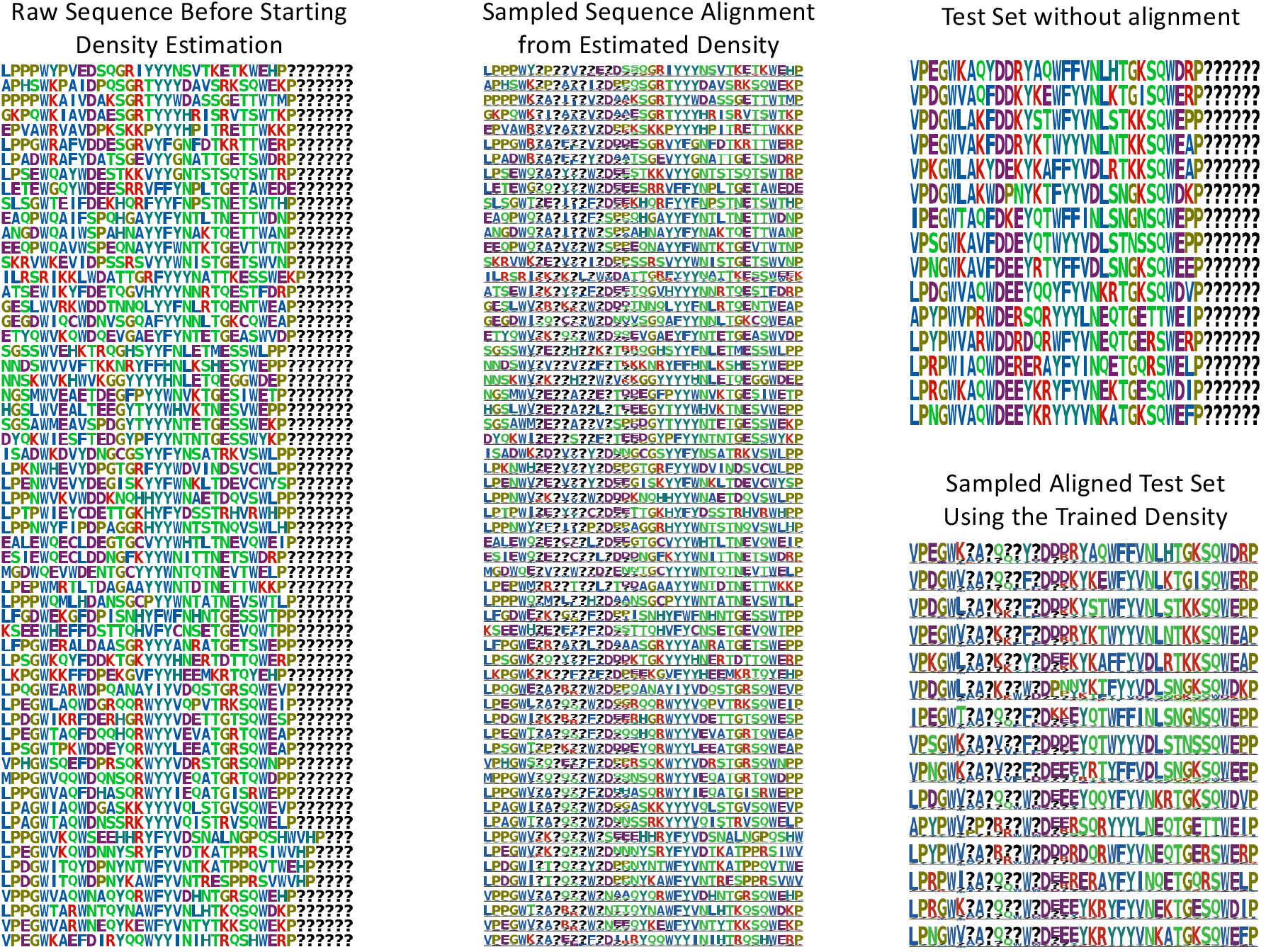
Generalization of sequence alignment on real data: The figure displays four plots showing the status of proteins associated with the WW domain. The plots show the unaligned sequences used to train the graphical model, the alignment of these sequences after density estimation, and the unaligned test sequences alongside their inferred alignments.

As shown in Figure 5, the model demonstrates robustness in both alignment and generalisation to novel, long sequences. This suggests that the inclusion of the stochastic network, modelled as a latent variable in the ELBO derivation, effectively captures the expressive features necessary for accurate alignment and representation.

It should be noted that the prior used was a VAE trained on 224 aligned sequences from the WW domain. On the basis of the results, it can be suggested that the model uses this prior as basic guidance to infer the optimal transformation to produce alignments that are similar to the reference information of the prior. This opens up the possibility of using VAEs on pre-existing sequence alignments as informative priors to recycle them and generate alignments based on a reference framework. However, the effectiveness of this alignment guide will depend on the quality of the training over the prior. Nevertheless, this approach has significant potential for the development of reference-guided alignments.

An important observation regarding Figures 2.D and 6.B is that, based on the results, it can be suggested that the light attention block Ψ is extracting sufficiently high-quality features for *z*_*P*_ to learn the transformation parameters, leading to accurate and precise alignments for this particular example. Furthermore, the generalization of this alignment to new sequences indicates that the distribution learned by *z*_*P*_ preserves the geometric transformation profiles specific to this protein family. By sampling from this distribution, the geometric structure of the features can be conserved across the family. This pattern recognition is analogous to the profile models used in Hidden Markov Models (HMM), such as HMMER [27], and may offer potential for future modelling approaches.

## Conclusion and Outlooks

As a proof of concept, the model yields promising results in multiple sequence alignment and generalisation through deep generative modelling. However, further testing and validation in specific applications such as mutation effect prediction and other protein engineering tasks are essential. In addition, it is important to assess the behaviour of the graphical model when it is estimated from scratch, as the current approach relies on an informative prior and introduces a new latent variable to improve feature extraction and inference of transformation parameters for long-range dependencies. While the model itself remains acyclic in this setting, the full end-to-end estimation procedure will have cyclic dependencies, which complicates parameter estimation in the model. We leave a further exploration of this matter for future work.

## Acknowledgement

This project has received funding from the European Union’s Horizon 2020 research and innovation programme under the Marie Sklodowska-Curie grant agreement No 801199, the Novo Nordisk Foundation through the MLSS Center (Basic Machine Learning Research in Life Science, NNF20OC0062606), and the Pioneer Centre for AI (DNRF grant number P1).

## A Appendix A

### A.1 Multi-output Gaussian Processes

A standard Gaussian process (GP) is a stochastic process where any finite set of random variables follows a multivariate normal distribution. This tool is very versatile and can be used for tasks related to non-linear regression. Formally, a GP is defined as:

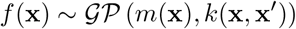

Where **x** is the input vector, *f*(**x**) is the random variable associated with each input **x, m** (**x**) = 𝔼[*f* (**x**)] is the mean function, representing the expected value of the process at **x** and *k*(**x, x**′) = 𝔼[(f (**x**) − **m** (**x**))(f (**x**′) − **m** (**x**′))] is the covariance function defined in terms of kernel function, which measures the covariance between the function values at **x**. Likewise, For any set of inputs {**x**_1_, **x**_2_, …, **x**_*n*_}, as well as the association with their corresponding output values {f (**x**_1_), *f* (**x**_2_), …, *f* (**x**_n_)} follow a joint multivariate normal distribution

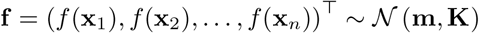

In view of the above, we can say that Multi-Output Gaussian Process (MO-GP) is a generalization of the standard GP to model a number of correlated outputs, with the aim of predicting several outputs at the same time. We can think of MO-GPs as a collection of GPs, each aiming to model a different output. We start from the assumption that each GP uses the same type of kernel function, but with different parameters focused on each task. Following the definition and notation outlined by [28], we can define the MO-GP as follows:

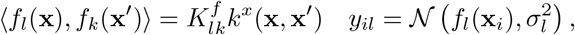

Here, *K*^*f*^ denotes a positive semidefinite (PSD) matrix representing the similarity between tasks, *k*^*x*^ is the covariance function over inputs, and 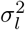 is the noise variance for the *l*^*th*^ task/channel. Similarly, for inference:

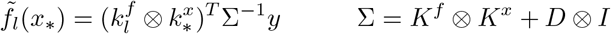

where ⊗ is the Kronecker product, 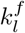 is the cofactor that selects the column *l*^*th*^ of *K*^*f*^, 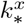 is the vector of covariances between the test sample *x*_*_ and the training points, *K*^*x*^ is the covariance matrix between all pairs of training points, *D* is a diagonal matrix in which the element (l, l)^*th*^ is the noise variance on that channel, and Σ denotes the global covariance matrix [28]. The simplest approach to extending MO-GPs is to model each output independently with single-output GPs, i.e. independent sets of GPs. This approach requires that the joint normal distribution over the output vector *y* is block diagonal with respect to the tasks or channels. Such a configuration prevents observations from one task or channel from influencing predictions for other tasks or channels [28], [29].

### A.2 Light Attention

Light Attention is a deep neural network architecture inspired by attention mechanisms in transformer networks. However, unlike traditional attention methods, it captures regions of interest through operations within convolutional networks, specifically using the Hadamard product. This architecture has primarily been employed to enhance representation and feature extraction in protein language models, such as ESM and ProtTrans, functioning as a pooling operator for downstream tasks, and has shown success in applications like subcellular localization prediction [26].

Originally designed as a pooling mechanism for protein language models, the expressive capabilities of Light Attention can be leveraged to address issues related to long-range dependencies, such as flat optimization landscapes when modeling long sequences. In contrast to the original implementation, where both convolutional layers process the same signal (see Figure 6.A), we made a slight modification to the approach by using two input sources instead: one convolutional layer handles the raw input, while the other extracts features from a distribution in the graphical model (see 6.B). This approach simulates what a cross-attention mechanism does on a smaller scale while preserving computational efficiency and speed. The light attention layer is denoted as 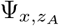 in the graphical model (refer to section A.2.1 and Figure 2.B), with the methodological scheme illustrated in Figure 6.

**Figure 6:**
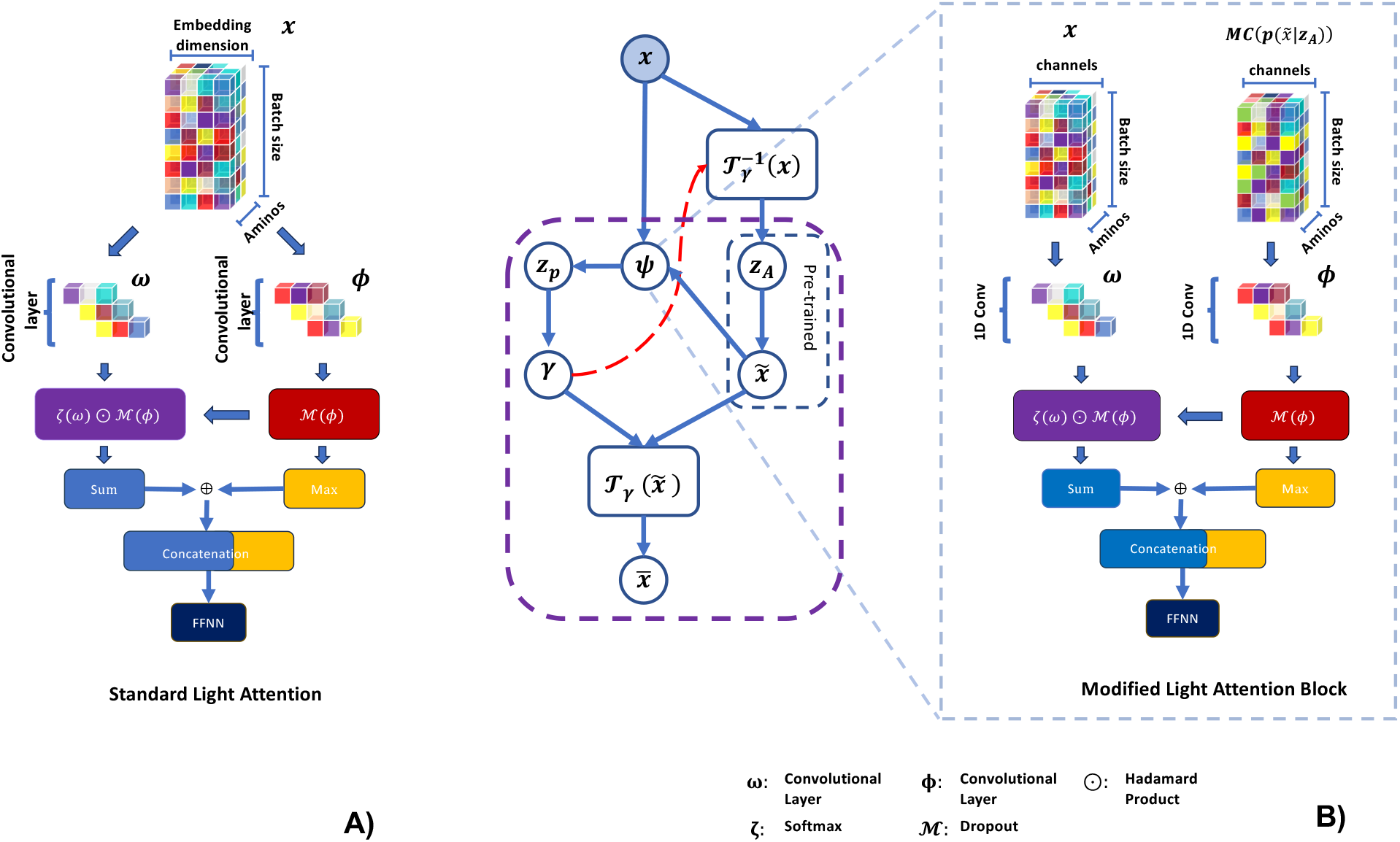
Light Attention building blocks: **A** illustrates the structure and functional blocks of light attention. First, given an input or embedding *x*, it passes through the convolutional layers *ω* and Φ to extract their respective features. The features obtained from Φ undergo a dropout, followed by a pointwise multiplication of the softmax of the features from *ω* and the resulting dropout from Φ. Additionally, pooling is applied to both the pointwise multiplication and the dropout of Φ. Finally, the features are concatenated and passed to a multilayer perceptron (MLP) to obtain the final features. **B** illustrates the integration of light attention as a latent variable, modelled as a normal distribution. Unlike the original light attention framework, this modification uses two inputs, each connected to different convolutional layers: x is connected to *ω*, while the Monte Carlo sample from the prior, defined as the pre-trained model of *z*_*A*_, is connected to 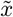 and then to Φ. The process then follows the same procedure as in the original implementation.

#### A.2.1 ELBO Derivation from the Model to Deal with Long Range Dependencies without Prior

In the main paper, we use an informative prior to guide the generalisation of sequence alignments. However, this raises the question of the implications of not having such a prior and starting the process from scratch. To explore this, we can attempt to derive the expression for ELBO under the following initial conditions:

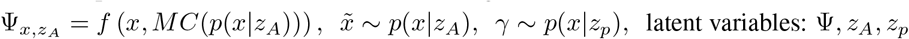

The joint distribution is defined as:

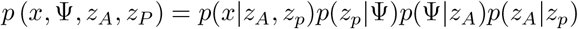

The evidence is defined as follows.

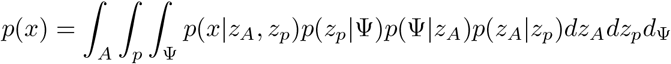

To derive log *p*(x), we get the following:

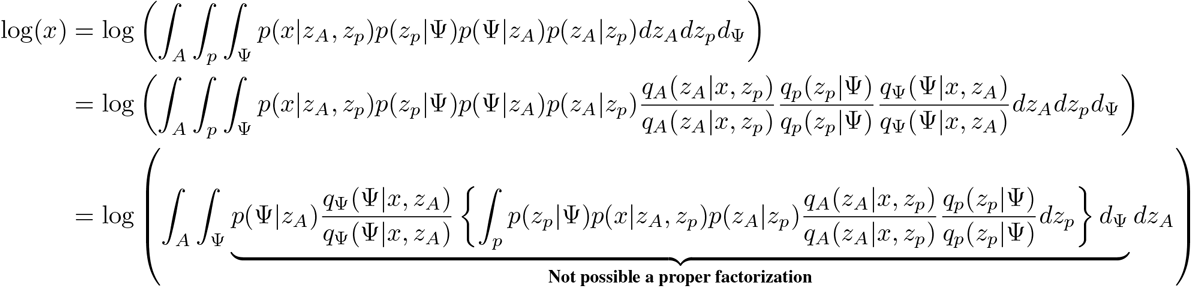

A detailed examination reveals that the inclusion of 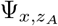 in the graphical model introduces a significant trade-off: the presence of a cycle within what was originally a Directed Acyclic Graph (DAG). This modification does not allow the derivation of a closed form expression for the ELBO which make it non-tractable. This limitation can be potentially be overcome using Markov Chain Monte Carlo (MCMC) techniques such as Gibbs sampling or Hamiltonian Monte Carlo, but we leave such considerations for future work.

